# Single-cell analysis of antiviral neuroinflammatory responses in the mouse dorsal raphe nucleus

**DOI:** 10.1101/780205

**Authors:** Kee Wui Huang, Bernardo L. Sabatini

## Abstract

Neuroinflammatory processes have been implicated in neurodegenerative and psychiatric diseases, and limit the utility of viruses for gene delivery. Here we analyzed 60,212 single-cell RNA profiles to assess both global and cell type-specific transcriptional responses in the mouse dorsal raphe nucleus following axonal infection of neurons by rabies viruses. We identified several leukocyte populations, which infiltrate the brain, that are distinct from resident immune cells. Additionally, we uncovered transcriptionally distinct states of microglia along an activation trajectory that may serve different functions, ranging from surveillance to antigen presentation and cytokine secretion. Our study also provides a critical evaluation of the compatibility between rabies-mediated connectivity mapping and single-cell transcriptional profiling. These findings provide additional insights into the distinct contributions of various cell types in the antiviral response, and will serve as a resource for the design of strategies to circumvent immune responses to improve the efficacy of viral gene delivery.

## INTRODUCTION

Neuroinflammatory processes are increasingly recognized for their importance in the etiology of neurological and psychiatric disorders. Recent studies have found significant associations between genes with known immune functions and diseases that include Alzheimer’s Disease and schizophrenia^1–4^. Microglia, the predominant resident immune cells of the central nervous system (CNS), express many of these disease-associated genes, and undergo transcriptional changes and diversification in both acute and chronic models of inflammation, aging, and neurodegeneration^5–8^. Transcriptional changes microglia during aging are further accompanied by increased lymphocyte infiltration in the brain^9^. However, the functions of these transcriptionally distinct microglial subsets and the interactions that trigger these changes during microglial activation have yet to be elucidated.

Immune responses limit the use of viruses as gene delivery vectors in the CNS. There has been great interest in the development and use of viral vectors for their applications in clinical gene therapy, including recently engineered variants of adeno-associated viruses (AAVs) that cross blood-brain barrier with high efficiency^10^. A wide variety of other viruses have been exploited as tools that have been essential for advances in basic neuroscience research, including single-stranded RNA viruses such as vesicular stomatitis virus (VSV), and rabies virus (RbV)^11–13^. The utility of many viruses are limited by their toxicity in neurons, as well as the clearance of both virions and virus-infected cells by the immune system. Understanding the various mechanisms by which the immune system and CNS responds to these viruses may facilitate the design of improved tools and strategies to circumvent these caveats.

Here we used high-throughput single-cell RNA sequencing (scRNA-seq) and assessed changes in gene expression and cell type composition of the mouse dorsal raphe nucleus (DRN) following infection of DRN projection neurons by glycoprotein-deleted rabies viruses (RbVs). Responses in the DRN were studied since pro-inflammatory signaling and immunological perturbations in the DRN have been implicated in behavioral and mood disorders that include from impulsivity, major depressive disorder, and bipolar disorder^14–18^. Analysis of single-cell transcriptomes from both RbV-injected and uninjected control mice revealed several types of infiltrating leukocytes in the DRN that are recruited by chemokines released from glial cell types. Analysis of the transcriptional changes by cell type also revealed both global and cell type-specific gene sets and pathways that underlie the antiviral defense response. Additionally, we identified transcriptionally distinct subsets of microglia, some of which minimally express canonical microglial genes, that may represent different functional states along an activation trajectory. Our results provide further insights into the shared and unique functions performed by various CNS cell types to mediate different facets of the immunological response.

## RESULTS

### Recruitment of circulating leukocytes into the DRN following rabies virus infection

Inflammatory responses were induced in the DRN by axonal infection of DRN neurons using glycoprotein-deleted RbVs of the SADΔG B19 strain, which are frequently used as tools for retrograde tracing to map neural connectivity^11, 13^. Retrograde axonal infection of DRN neurons by RbVs permitted the physical separation of the injection site from the location of the infected neuronal cell bodies, and reduced the effects of physical tissue damage.

Tissue containing the DRN was collected from RbV-injected animals (4 mice – 2 male, 2 female) 7 days post-injection. RbVs were injected into a pair of brain regions that were both innervated by DRN neurons, and included the striatum (Str), dorsal lateral geniculate nucleus (dLGN), nucleus accumbens (NAc), and substantia nigra (SN) (Fig. 1a). Tissue chunks were dissociated into live whole-cell suspensions, and single-cell RNA-seq (scRNA-seq) libraries were prepared using the inDrop v3 platform^19, 20^. Inhibitors of neural spiking activity, transcription, and translation were included to reduce the effects of tissue dissociation on gene expression^21^. Cells from RbV-injected animals were analyzed together with cells collected from uninjected animals, and datasets were merged using CCA-based dataset alignment methods^22^. Low-quality cells and putative multiplets were manually identified and discarded prior to analysis of differential gene expression (see Methods). Our final merged dataset contained a total of 60,212 cells: 20,581 cells in the RbV group (10,065 male, 10,516 female), and 39,631 cells in the Control group (17,496 male, 22,135 female). The Control dataset included cells previous analyzed and described in a separate study^23^. Cells were sequenced to a mean read depth of 62,061 reads/cell (min. = 20,001; median = 47,823; IQR = 43,294; max. = 870,064), 2,603 UMI-filtered mapped reads (UMIFMs) (min = 501; median = 2,093; IQR = 1,849; max. = 17,997) and mean gene detection rate of 1,027 genes/cell (min. = 201; median = 896; IQR = 706; max. = 5,518). Separation of the cells by condition showed that each group was sequenced to comparable read depths with the RbV group having a higher mean (Control: 1^st^ quartile = 30,670, median = 45,709, mean = 58,359, 3^rd^ quartile = 69,928; RbV 1^st^ quartile = 34,090, median = 52,542, mean = 69,191, 3^rd^ quartile = 85,428). However, the UMIFM count (Control 1^st^ quartile = 1,574, median = 2,338, mean = 2,856, 3^rd^ quartile = 3,499; RbV 1^st^ quartile = 1,082, median = 1,649, mean = 2,117, 3^rd^ quartile = 2,605) and gene detection rates (Control 1^st^ quartile = 741, median = 1,041, mean = 1,173, 3^rd^ quartile = 1,446; RbV 1^st^ quartile = 420, median = 612, mean = 745.8, 3^rd^ quartile = 942) were slightly lower in the RbV group compared to the Control group.

**Fig. 1:**
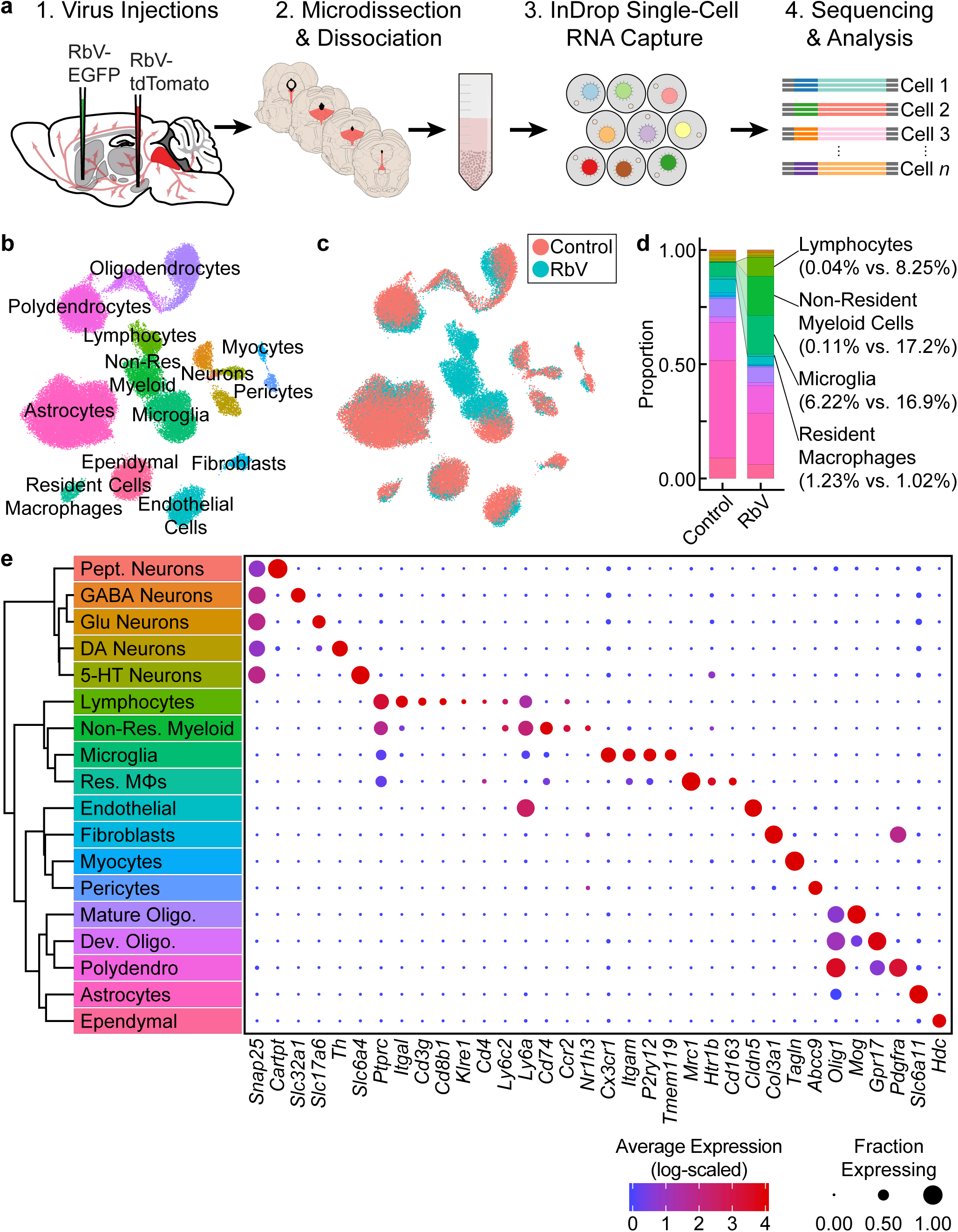
Single-cell transcriptional analysis of responses to viral infection in the brain. **(a)** Experiment schematic. Unpseudotyped SADΔG B19 rabies viruses (RbV) were injected into 8-10 week old C57BL/6J mice. Each animal received a pair of injections into two different regions innervated by DRN 5-HT neurons. Tissue containing the DRN was dissected 7 days post-injection and dissociated into whole cell suspensions. scRNA-seq libraries were generated using the microfluidic-based inDrop platform. Age-matched uninjected animals were used as the Control group. **(b)** UMAP plot of merged dataset containing 60,212 cells. Control and RbV datasets were merged using CCA-based dataset alignment methods. Individual points representing single cells are color-coded by cell class/type as shown in (**e**). **(c)** UMAP plot of merged dataset with cells color-coded by experimental condition (RbV or Control). **(d)** Stacked bar plot showing the relative proportion of each cell type in RbV and Control groups. Cell class/type categories are color-coded following the same scheme as (**b**). **(e)** *Left*: Dendrogram with cell class/type labels corresponding to the cluster labels in (**b**). *Right*: Dot plot showing expression of example genes (columns) used to identify the major cell classes/types (rows). The color of each dot represents the average log-scaled expression of each gene across all cells in a given cluster, and the size of the dot represents the fraction of cells in the cluster in which transcripts for that gene were detected.

Cells in the merged dataset were clustered in a CCA-aligned space (see Methods). Inspection of genes enriched in each cell cluster showed that all of the resident cell types that we previously identified in the reference dataset^23^ were present in the RbV group (Fig. 1b, e). However, there was a significant expansion in the proportion of immune cells in the RbV group (Fig. 1c–d). This included an increase in the proportion of microglia (*n* = 5,949 cells, 16.9% of RbV vs. 6.2% of Control), but not resident macrophages (Res. MΦs) (*n* = 698 cells, 1.02% RbV vs. 1.23% of Control). We also identified both myeloid and lymphoid cell clusters that were not previously found in the uninjected Control group. A large cluster of myeloid cells that are distinct from microglia and resident MΦs formed a significant proportion of the cells in the RbV-injected group (*n* = 3,575 cells, 17.2% of RbV vs. 0.11% of Control), and the proportion of lymphocytes was also greatly increased in the RbV-injected group (*n* = 1,714 cells, 8.25% of RbV vs. 0.04% of Control). The appearance of these non-resident immune cells is likely due to the infiltration of the brain parenchyma by circulating leukocytes, since blood was removed from the brain via transcardial perfusion prior to tissue collection. A comparison of the proportion of immune cells in groups sorted by RbV injection sites suggested that leukocyte infiltration scales with infection magnitude, which was assessed as the mean number of RbV-infected neurons labeled by each injection (Supplementary Fig. 1a–e). While samples collected from mice that received a pair of RbV injections into the Str and dLGN showed an increase in the proportion of microglia compared to Control/Uninjected animals (Uninjected = 6.2%; Str-dLGN = 16.3%; NAc-SN = 17.5%), animals that received injections into SN had a larger increase in the proportion of lymphocytes (Uninjected = 0.04%; Str-dLGN = 0.75%; NAc-SN = 14.5%) and non-resident myeloid cells (Uninjected = 0.11%; Str-dLGN = 0.55%; NAc-SN = 30.9%) (Supplementary Fig. 1a–b). Injection of RbV into SN infected an order of magnitude more neurons than Str, NAc, or dLGN injections (Supplementary Fig. 1c–e), suggesting that the number of infected cells correlates with the magnitude of the immune response and leukocyte recruitment.

### Rabies virus transcripts are detected in both neurons and microglia

To identify projection neurons that are infected by RbVs, we calculated RbV gene set expression scores for each cell (see Methods) and examined the distribution of RbV gene transcripts in each cell type in the RbV group (Supplementary Fig. 2). As expected from the innervation of the injection sites by 5-HT neurons, the 5-HT neuron cluster (*n =* 275 cells in RbV group) had the highest average RbV expression scores. However, the rate of detecting cells containing RbV transcripts was low. The low yield of RbV-infected neurons could be due to the following: (i) since cells were not sorted during tissue dissociation, only a small fraction of RbV-infected neurons were captured during the cell encapsulation and mRNA capture given the typical capture efficiency of 30-50%; (ii) RbV-infected neurons may have poor survival during the tissue digestion and dissociation and were therefore relatively depleted from the whole cell suspensions; (iii) given the relatively low read depths and the lack of enrichment for RbV transcripts during library preparation, there are likely to be transcripts from RbV-infected cells that were not sequenced (drop-outs in scRNA-seq); (iv) few RbV-infected neurons were labeled due to the low labeling efficiency of the viruses. Despite the low detection rate, the RbV-infected cells that were identified retained sufficient transcriptional information for clustering and assignment of cell class/type identity.

RbV transcripts were also detected in non-neuronal cells: microglia (*n =* 3,483 cells in RbV group), non-resident myeloid cells (*n =* 3,530 cells in RbV group), and astrocytes (*n =* 4,605 cells in RbV group) showed the next 3 highest average RbV expression scores and had more than 1 cell above the score threshold (red line in Supplementary Fig. 2a). These cells are unlikely to have been directly infected by RbV, given the tropism of RbVs, the lack of transsynaptic spread by the glycoprotein-deleted RbVs used, and the long distance of the DRN from the injection sites. A comparison of RbV transcript counts between 5-HT neurons, predicted to be the cells directly infected by RbVs, and the other 3 non-neuronal cell types showed that maximum number of RbV transcript counts in these non-neuronal cells was at least an order of magnitude lower than 5-HT neurons (Supplementary Fig. 2b). We hypothesize that the detection of RbV transcripts in these cells are due to their phagocytosis of mRNA-containing material released from infected neurons. The non-zero background counts of RbV transcripts that we observed in other cells may also be due to the capture of free-floating RbV transcripts released from the physical disruption of some RbV-infected cells during tissue digestion and dissociation.

RNA viruses can be recognized via pattern recognition receptors that include RIG-I and RIG-I-like receptors (e.g. MDA5), which detect foreign RNA. Since RIG-I and RIG-I-like pathways are thought to be the primary means of detecting infection by RNA viruses, we calculated the per cell expression score of genes in the KEGG *“RIG-I-like signaling pathway”* gene set (Supplementary Fig. 2c). All neuron cell types had low scores for expression of genes involved in RIG-I-like signaling. Surprisingly, we did not observe a positive correlation between the RIG-I signaling gene set expression score and the RbV gene set expression score (RbV group, all cells, Pearson R = 0.02) (Supplementary Fig. 2d), whereas a positive correlation was observed between expression of different RbV genes despite the occurrence of drop outs (RbV group, all cells, Pearson R = 0.57) (Supplementary Fig. 2f). Genes downstream of RIG-I, such as *Tmem173* (STING) were also low even in 5-HT neurons with high expression of RbV transcripts (Supplementary Fig. 2e). However, RbV components, such as the P protein, which are expressed by the glycoprotein-deleted mutants we used may inhibit interferon signaling in infected neurons^24–26^. The low expression of interferon stimulated genes (e.g. *Isg15*) and low RIG-I-like signaling gene set expression scores across neuronal types may also indicate an underlying difference in the function of neurons versus glia in innate immunity – astrocytes and microglia, which form close associations with synapses and neurons, may serve more prominent roles in the detection of pathogens and neuronal infection due to the suppression of these pathways in neurons.

### Identification of global and cell type-specific transcriptional changes

To assess the population-level transcriptional responses to RbV infection, we first performed differential expression (DE) tests on simulated “bulk” RNA-seq samples separated into RbV and Control groups (Fig. 2a). “Bulk” samples were simulated by averaging UMI counts for each gene across all cells in a group regardless of cell type. Many genes involved in antiviral immune responses that were expressed at low levels in the Control group were strongly up-regulated in the RbV group. These included interferon response genes such as *Isg15,* major histocompatibility complex (MHC) genes such as *H2-Aa* and *H2-D1*, and genes that are highly expressed by infiltrating leukocytes such as *Ptprc*. Gene set enrichment analysis (GSEA) using the *MSigDB* Hallmark gene sets^27, 28^ showed that genes involved in type I and type II interferon responses were highly up-regulated, as well as various signaling pathways such as complement, IL-2, IL-6, and TNFα (Fig. 2b). Genes involved in cell division were also up-regulated, consistent with the proliferation and clonal expansion of microglia and T cells upon activation. Our results are also consistent with prior studies that have used bulk tissue profiling methods to evaluate transcriptional changes in the CNS following exposure to RbVs^29–31^.

**Fig. 2:**
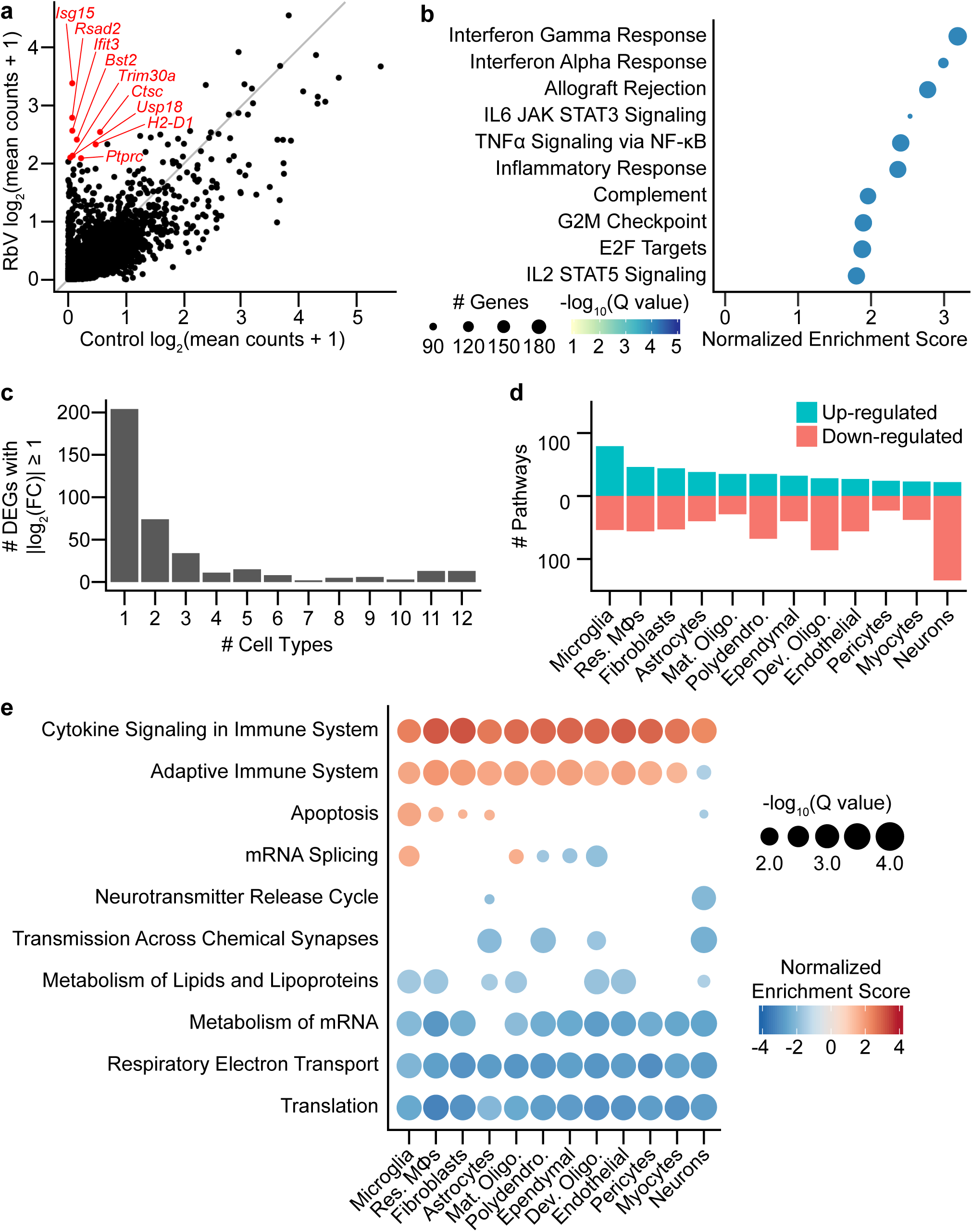
Rabies infection induces both global and cell type-specific transcriptional changes. **(a)** Scatter plot comparing averaged expression of each gene in simulated “bulk” RNA-seq for RbV and Control groups. Several of the genes most enriched in the RbV group are labeled in red. Gray line indicates the line of unity (*x* = *y*). **(b)** Dot plot of the top *MSigDB* hallmark gene sets that were significantly enriched and up-regulated in the RbV group (5% FDR, Benjamini-Hochberg correction, Normalized Enrichment Score > 0). **(c)** Bar plot showing the number of differentially expressed genes (DEGs) that were significantly changed (388 DEGs with Q value < 0.01 and |log_2_ fold-change| > 1). DE genes were either a part of global response programs (35 genes significant in ≥ 9 resident cell types) or cell type-specific responses (204 genes significant in 1 of 12 resident cell types). **(d)** Number of Reactome pathways that are either up-regulated (blue) or down-regulated (red) in the RbV group compared to controls. **(e)** Dot plot showing Reactome pathways that are altered in each type of resident cell. The size of each dot represents the number of genes in each cell type (columns), and the color of each dot indicates the normalized enrichment score of each pathway (rows). Pathways that were not significantly enriched (Q value ≥ 0.05, Benjamini-Hochberg correction) are not displayed.

We assessed the transcriptional changes in resident cell types by performing DE tests and gene set enrichment analysis (GSEA) separately for each cell type/class. Most differentially expressed genes (DEGs) that were strongly up- or down-regulated (388 genes with Q value < 0.01, absolute log_2_ fold-change > 1) were found to be differentially expressed in only 1 of the 12 resident cell types (Fig. 2c), and were considered cell type-specific DEGs (204 of 388 genes). Several genes were differentially expressed in 9 or more cell types and were considered part of a “global” response program (35 of 388 genes). These genes included many of the interferon-stimulated genes such as *Isg15* and other genes involved in the inflammatory response. Pathway analysis using GSEA on Reactome pathway gene sets showed that the cell types varied in their responses. Neurons had the most down-regulated pathways (28 up-regulated, 148 down-regulated), whereas microglia had the highest number of up-regulated pathways (96 up-regulated, 61 down-regulated) (Fig. 2d). Reactome pathways that were globally up-regulated included “*Cytokine Signaling in Immune System”* and “*Adaptive Immune System”*, whereas globally down-regulated pathways were mostly related to metabolism, such as *“Translation*”, *“Respiratory Electron Transport”*, *“Metabolism of Lipids and Lipoproteins”*, and *“Metabolism of mRNA”* (Fig. 2e). The down-regulation of these pathways may be driven by the antiviral response in an attempt to inhibit viral replication and spread, and may underlie the reduced UMIFM counts and gene detection rate in the RbV group relative to the Control group.

Several pathways were enriched in only a subset of cell types but not found to be cell type-specific. Genes in the *“Apoptosis”* set were up-regulated in resident immune cells, fibroblasts, and astrocytes, but were down-regulated in neurons. The *“mRNA Splicing”* gene set was also up-regulated in microglia and mature oligodendrocytes while being down-regulated in other glial cell types including developing oligodendrocytes. Genes involved in neurotransmission were down-regulated in neurons and glial cell types that express neurotransmitter receptors or are involved in the regulation of synaptic transmission, including astrocytes and polydendrocytes. A reduction in neurotransmission may contribute to the host defense by limiting the spread of neurotropic viruses across synapses, and has been suggested to be induced by IFNγ signaling^32^.

To identify the cell types and signaling molecules mediating the recruitment of infiltrating leukocytes, we sorted DE genes based on their gene ontology annotations to find differentially expressed cytokines and chemokines. Microglia and resident MΦs showed the highest increase in cytokine expression compared to other resident cell types (Supplementary Fig. 3a). Cell types of the neurovascular unit, which include endothelial cells, astrocytes, pericytes, and fibroblasts/fibroblast-like cells, and ependymal cells at the interface with the ventricular system showed the next highest increase in cytokine production. Neurons showed the least increase in cytokine expression. Several pro-inflammatory chemokines, such as *Cxcl9*, *Cxcl10*, *Ccl2*, *Ccl5*, and *Ccl7*, were released by multiple cell types. Most cell types in the RbV group released a distinct set of cytokines (Supplementary Fig. 3b), with fibroblasts expressing the largest set cytokines. In contrast to a recent study using a “viral déjà vu” model^33^, we did not detect *Ccl2* expression in neurons. We speculate that this may be due to a difference in the models used – our study examines changes that occur during the primary responses on first encounter with the virus, whereas the “viral déjà vu” model investigates secondary responses.

### Infiltrating leukocytes are transcriptionally diverse

To identify the types of immune cells involved in the response to RbV infection, we performed subclustering on the immune cell subset (Fig. 3a, Supplementary Fig. 4a). Infiltrating leukocytes were separated into two main groups that were of either myeloid or lymphoid lineage. Iterative subclustering resolved the lymphoid cell cluster into at least 3 distinct types. T cells (*Cd3g*, *Cd3e*) were comprised of both CD8^+^ effector T cells (*Cd8a*, *Cd8b1*, *Tcf7*, *Prf1*, *Gzma*), and a smaller number of CD4^+^ helper T cells (*Cd4*, *Il2ra*, *Ctla4*). Natural killer (NK) cells were also present in the lymphoid group and were identified by their expression of genes such as *Klre1*, *Prf1*, and *Gzma*. Non-resident myeloid cells were transcriptionally heterogeneous, and were comprised of several populations that were distinct from both of the resident myeloid cell types (Supplementary Fig. 4a–i). The majority of non-resident myeloid cells were monocytes (*Ccr2*, *Fn1*, *Plac8*, *Lyz2*, *Ly6c2*^lo/mid^) and monocyte-derived macrophages (moMΦs) (*Ly6c2*^hi^, *Nr1h3*, *Ly6a*, *H2-Aa*, *Ms4a6d*, *Cd74*). Several distinct clusters of dendritic cells (DCs) were also identified, which included monocyte-derived dendritic cells (moDCs) (*Il1b*, *Il1dr1*, *H2-Aa*, *Cd74*, *Ifitm1*), conventional dendritic cells (cDCs) (*Ccr7*, *Il4i1*, *Cd74*, *Cacnb3*), and plasmacytoid dendritic cells (pDCs) (*Ly6d*, *Siglech*, *Irf8*, *Runx2*).

**Fig. 3:**
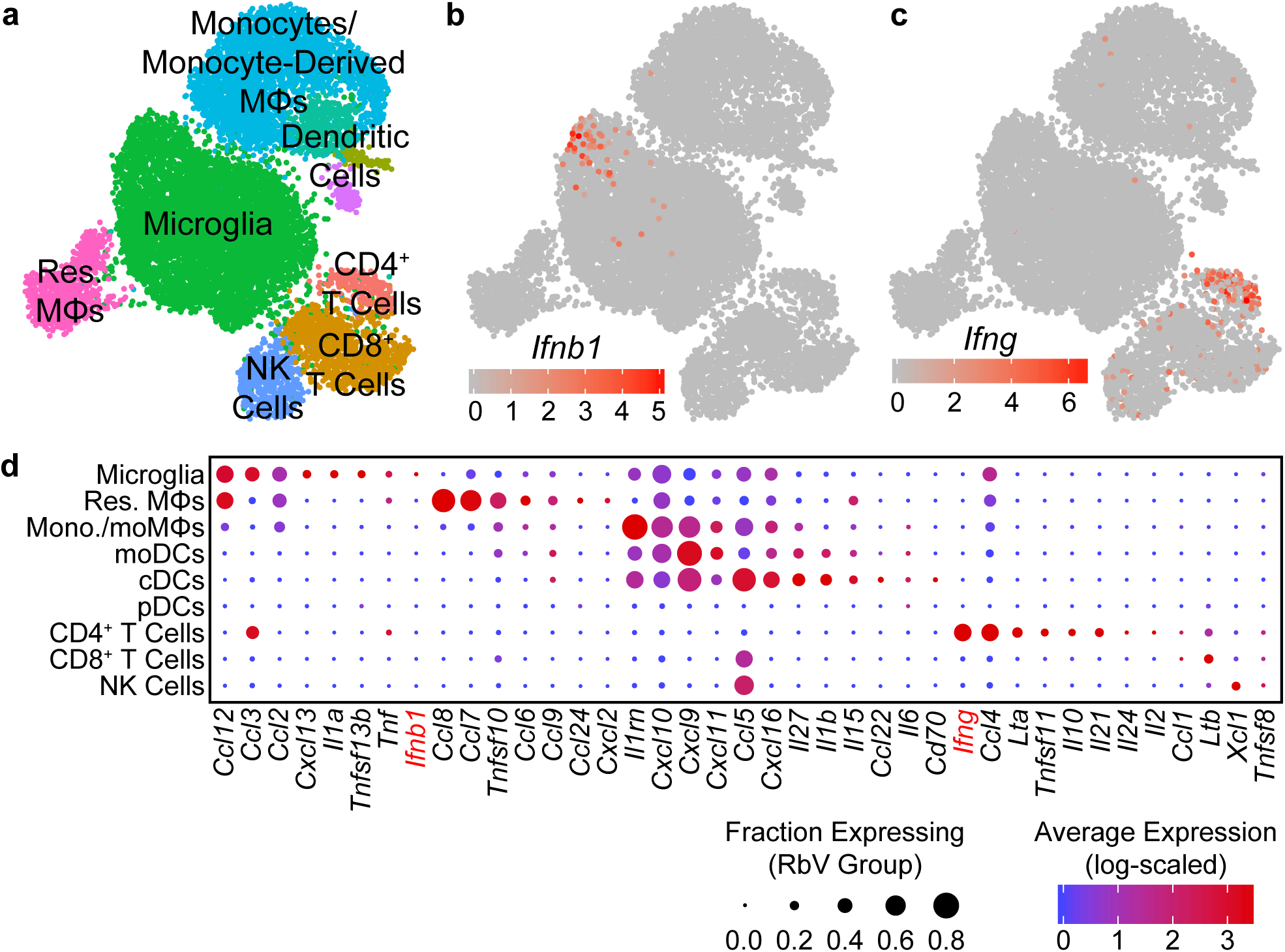
Type I interferon responses are induced by IFNβ produced in microglia. **(a)** UMAP plot of the immune cell subset. Individual points represent single cells, which are color coded by their assigned cell type identity. **(b)** UMAP feature plot showing *Ifnb1* expression, in which the color of each cell represents the log-scaled UMI counts. *Ifnb1* transcripts are only detected in a small subset of microglia. **(c)** UMAP feature plot for *Ifng* expression. CD4+ T cells are the primary source of *Ifng*. **(d)** Dot plot showing the expression of selected cytokine genes (columns) in each immune cell type (rows). The dot color represents the average log-scaled expression of each gene in a given cell cluster, and the size of the dot represents the fraction of cells in that cluster in which transcripts for that gene were detected.

Since genes involved in immune processes are increasingly associated with Alzheimer’s Disease (AD), we assessed the expression of several AD-associated genes by cell type (Supplementary Fig. 4j). While several genes such as *App* were enriched in neurons, AD-associated genes involved in immune processes were expressed in specific cell types. *Trem2* was specifically expressed in microglia, whereas genes in the *Ms4a* family (e.g. *Ms4a7*, *Ms4a6c*) were primarily expressed by both resident MΦs and infiltrating moMΦs. *Ms4a4b* was also expressed in infiltrating lymphocytes. Resident MΦs also expressed high levels of *Apoe* in addition to astrocytes. Several AD-associated genes were also differentially expressed between the RbV and Control groups. We observed a decrease in *Trem2* and *Cd33* expression in response to RbV infection, whereas expression of MHC class I genes (*H2-Eb1*) and *Ms4a* genes were increased.

### Type I interferon responses are mediated by release of IFNβ from microglia

To identify the primary mediators of the antiviral transcriptional responses, we assessed expression of interferons in each cell type. Of the genes encoding type I or II interferons, only transcripts for *Ifnb1* and *Ifng* were detected in our dataset. *Ifnb1* expression was restricted to a small subset of microglia (Fig. 3b), while *Ifng* was expressed at high levels by CD4^+^ T cells, and at lower levels by CD8^+^ T cells (Fig. 3c). Surprisingly, genes for IFNα were not detected in any cells, despite the presence of pDCs that are typically the main source of type I interferons in the periphery^34^. These results are consistent with previous reports of IFNβ production from microglia to limit viral spread^35^. However, our results contrast with previous reports that identified astrocytes as the primary source of IFNβ^36^, which may be due to differences in the methods used to identify IFNβ-producing cells. Given the low rate in which RbV-infected cells were detected, we are unable to rule out expression of IFNα/β from infected neurons.

In addition to interferons, many other cytokines were also differentially expressed between immune cell types (Fig. 3d). The pro-inflammatory cytokine *Il1a* was expressed specifically in microglia, whereas *Il1b* was expressed in cDCs and moDCs. Infiltrating myeloid cells expressed high levels of several pro-inflammatory cytokines, including *Ccl5* and *Cxcl9*. Microglia, resident MΦs, and CD4^+^ T cells also expressed *Tnf*, which may facilitate leukocyte infiltration via its effects on neurovascular cells and the blood brain barrier^37, 38^. Other members of the TNF superfamily were also expressed in different cell types, including *Tnfsf10* in resident MΦs and monocytes/monocyte-derived cells, *Ltb* in T cells, and *Tnfsf11* in CD4^+^ T cells. Several anti-inflammatory cytokines were also expressed by specific cell types: *Il10* was expressed specifically in CD4^+^ T cells, whereas the IL-1 receptor antagonist gene *Il1rn* was expressed by microglia and several infiltrating myeloid cell types.

### Microglia occupy distinct states along an activation trajectory

Since microglia had the highest number of up-regulated DE genes and pathways in response to the RbV infection, we performed subclustering on the microglial subset to determine if there are transcriptionally distinct states or subtypes of activated microglia. Subclustering divided microglia into at least 7 distinct subclusters (Fig. 4a). Subclusters differed in their proportion of cells from groups separated by condition (RbV / Control) and injection sites (Uninjected / Str-dLGN / NAc-SN). Subclusters that had a higher proportion of cells from the Control group included subclusters I (77.0% in Uninjected; 12.5% in Str-dLGN; 10.4% in NAc-SN) and subcluster II (79.5% in Uninjected; 14.5% in Str-dLGN; 6.0% in NAc-SN). Subclusters with microglia primarily from the RbV group included subclusters V (1.0% in Uninjected; 22.4% in Str-dLGN; 76.6% in NAc-SN), VI (0.3% in Uninjected; 3.2% in Str-dLGN; 96.4% in NAc-SN), VII (0.5% in Uninjected; 12.0% in Str-dLGN; 87.5% in NAc-SN), and VIII (0.3% in Uninjected; 91.6% in Str-dLGN; 8.1% in NAc-SN). Subcluster III (13.2% in Uninjected; 69.8% in Str-dLGN; 17.0% in NAc-SN) and subcluster IV (59.3% in Uninjected; 6.5% in Str-dLGN; 34.2% in NAc-SN) showed intermediate proportions between RbV and Control groups.

**Fig. 4:**
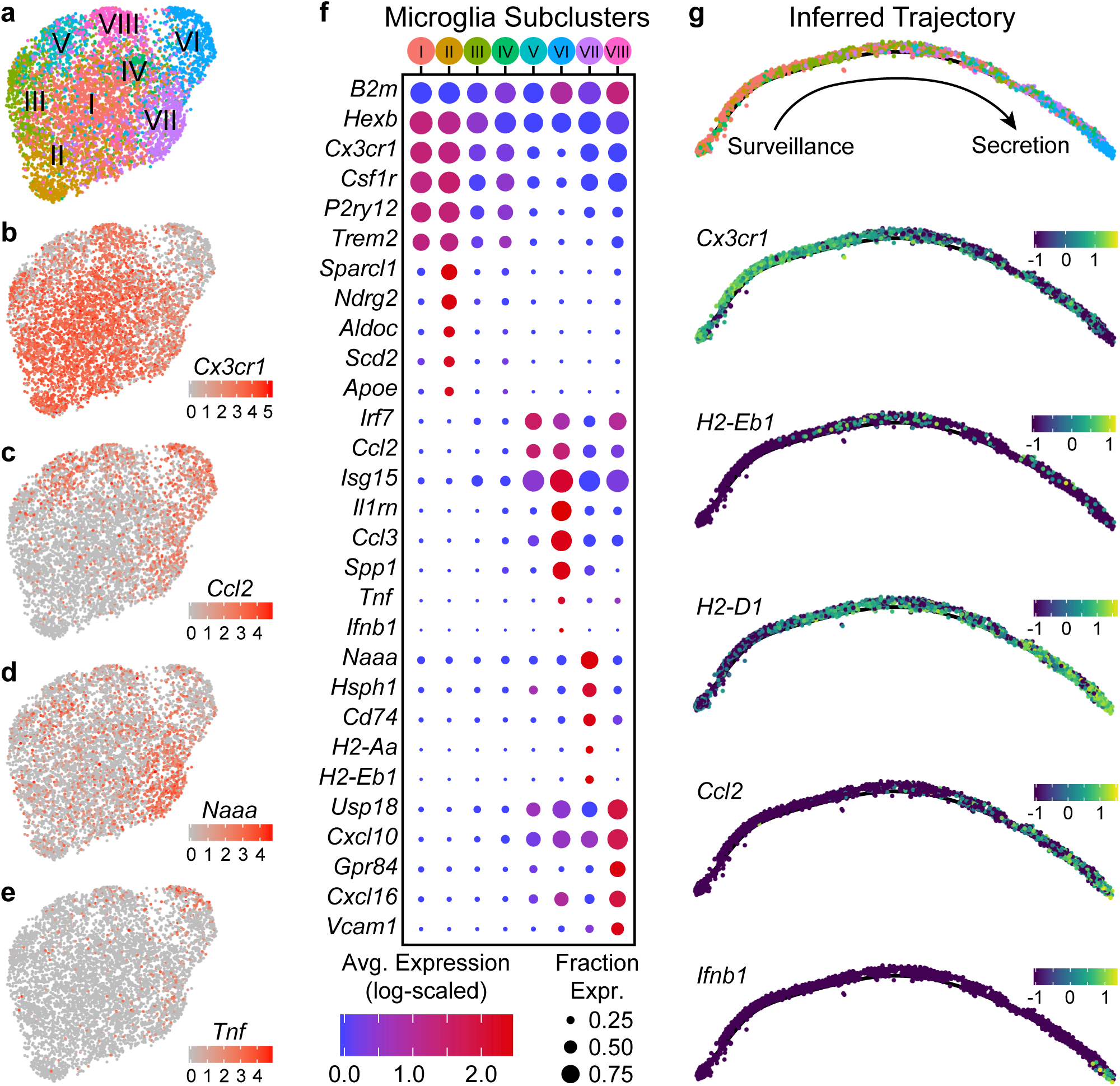
Microglia are found in distinct transcriptional states along an activation trajectory. **(a)** UMAP plot showing subclusters of microglia found using SNN graph-based clustering (see Methods). Points are single cells color-coded by subcluster assignment. **(b–e)** UMAP feature plots showing expression of genes that are differentially expressed between microglia subclusters. Cells are color-coded by their log-scaled UMI counts for the indicated gene. **(f)** Dot plot showing expression of several genes (rows) that are differentially expressed between microglia subclusters (columns – color-coded according to (**a**)). **(g)** Inferred trajectory of microglial activation. Genes that were differentially expressed between microglia subclusters were used for trajectory inference. Individual cells were arranged along a single unbranched trajectory based on their pseudotime values. *Top*: Cells are color-coded by subclusters, following the color scheme in (**a**). Cells from subclusters I-II are on the left-most end of the trajectory, while cells from subcluster VI are on the opposite end. The remaining subclusters were distributed along the middle of the inferred trajectory. *Middle/Bottom*: Inferred graphs with cells color-coded by their expression of the indicated genes, quantified as log_10_(counts+0.1), which are differentially expressed between microglia sub-clusters.

Differential expression tests between subclusters showed that they differ in expression of various genes that include genes that are typically used as “markers” for microglia such as *Cx3cr1*, *Csf1r*, *P2ry12*, and *Trem2* (Fig. 4b–f). Subclusters I and II had the highest expression of these canonical “marker” genes, and are likely to be microglia in a “resting” or surveillance state. Expression of these canonical “marker” genes was lower in the remaining subclusters and anti-correlated with expression of interferon-stimulated genes such as *Isg15* (all microglia, *Cx3cr1* vs. *Isg15*, Pearson R = -0.56). Subclusters VII, VIII, V, and VI express progressively lower levels of *Cx3cr1* (in the stated order), and instead expressed *Isg15*, *Irf7*, and *Ccl2* at higher levels. Between subclusters V-VIII genes involved in different immunological processes such as antigen presentation (e.g. *Cd74*, *H2-Aa*, *H2-Eb1*) and cytokine secretion (e.g. *Ifnb1*, *Tnf*) are differentially expressed, suggesting that these subclusters are functionally distinct.

To assess whether subclusters V-VIII reflect distinct states or subtypes of microglia, we used trajectory inference methods based on the genes that are differentially expressed between the subclusters^39, 40^. We hypothesized that branches in the inferred trajectory/graph occupied by different subclusters would suggest that these subclusters define distinct activation endpoints for microglia and diversification of activated microglia, whereas an unbranched trajectory with a single edge/path would instead suggest that the subclusters define discrete states along a continuous trajectory and describe the transitional states that can be occupied through several stages of microglial activation. The graph that was constructed from the microglia transcriptomic data suggested that the subclusters lie along an unbranched trajectory (Fig. 4g). Subclusters I and II that express high levels of *Cx3cr1* and *P2ry12* were enriched on one extreme of the inferred trajectory, and describe the “resting” or surveilling state of microglia. Subcluster VIII cells that are enriched for genes involved in antigen presentation, such as *Cd74* and *H2-Eb1*, were densest in the middle of the trajectory where there is also an increase in expression of interferon response genes such as *Isg15*. Subcluster VI cells, which are enriched in expression of cytokines such as *Ccl2*, *Tnf*, and *Ifnb1*, were found at the other extreme of the trajectory, and describe a secretory state. Given the higher abundance of infiltrating lymphocytes in the NAc-SN group (majority of subcluster VI microglia) compared to the Str-dLGN (majority of subcluster VIII microglia), we speculate that the transition of microglia from an antigen presentation state the middle of the inferred activation trajectory to the secretory state may be mediated by interactions between microglia and T cells.

### RbV infection alters the structure of intercellular interactions in the DRN

To assess changes in intercellular communication between specific cell types induced by the immune response, we used *CellPhoneDB* to predict significant interactions between the various cell types from our scRNA-seq data^41, 42^. Significant interactions were assessed for RbV and Control groups separately (Supplementary Fig. 5), and the difference between the two was used to infer changes in intercellular interactions (Fig. 5). Interactions among resident cell types under control conditions were highest between fibroblasts and cells of the neurovascular unit, although we anticipate that the number of interactions we infer here is likely to be an underestimate since the analysis package may not include interactions mediated by neurotransmitter release. A comparison of interactions between RbV and Control groups showed an overall decrease in intercellular communication between most resident cell types. Interactions with infiltrating cell types, such as monocytes and dendritic cells, showed a large increase as expected from their relative absence in the Control group. Among the resident cell types, microglia and resident MΦs had the highest increase in interactions. In particular, microglia increased in their interactions with CD4^+^ T cells, consistent with our hypothesis that microglia-T cell interactions may be involved in the progression of microglia along the activation trajectory. Microglia also increased in the number of inferred interactions with 5-HT neurons relative to the other neuron types. We hypothesize that this may be indicative of increased interactions between microglia and RbV-infected neurons.

**Fig. 5:**
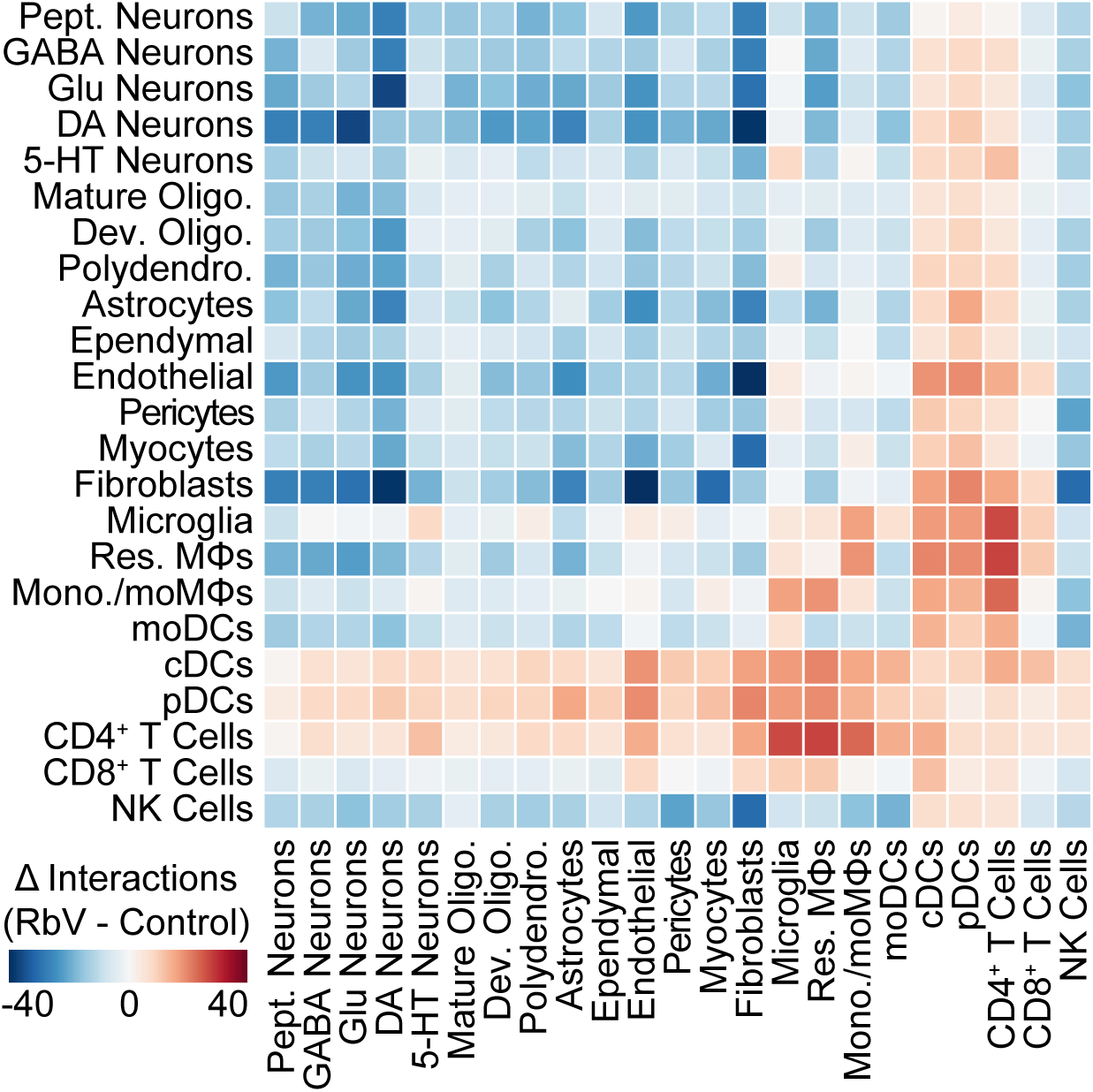
The structure of intercellular interactions is altered by the immune response. Heatmap showing the change (RbV - Control) in the number of interactions between cell types resulting from RbV infection. Interactions were inferred from single cell expression data using *cellphoneDB*. Red colors indicate an increase in the number of interactions between a pair of cell types in the RbV group relative to Control, whereas blue colors indicate a decrease in the number of interactions.

## DISCUSSION

Here we characterize both global and cell type-specific transcriptional responses in the CNS following neuronal infection by RbVs. We reveal the transcriptional diversity of both resident and infiltrating immune cells populations that mediate distinct aspects of the host antiviral defense mechanisms. These results suggest that microglia and infiltrating T cells serve key roles in orchestrating the antiviral response via Type I and Type II interferon signaling. Our data also shows that cytokine signaling induced by viral infection leads to the down-regulation of metabolic processes and neurotransmission, which may limit the spread of neurotropic viruses. We also describe transcriptionally distinct subsets of microglia that are likely to represent discrete transitional states along an activation trajectory during the progression of the immune response. Additionally, we outline the changes in cell type-specific intercellular interactions involved in different aspects of the multifaceted response, which led to our prediction that helper T cells may mediate the progression of microglia along an activation trajectory (Fig. 6). Our study provides additional insights into the distinct immunological functions of various cell types in the brain, and presents several testable models and hypotheses for experimental validation in future studies.

**Fig. 6:**
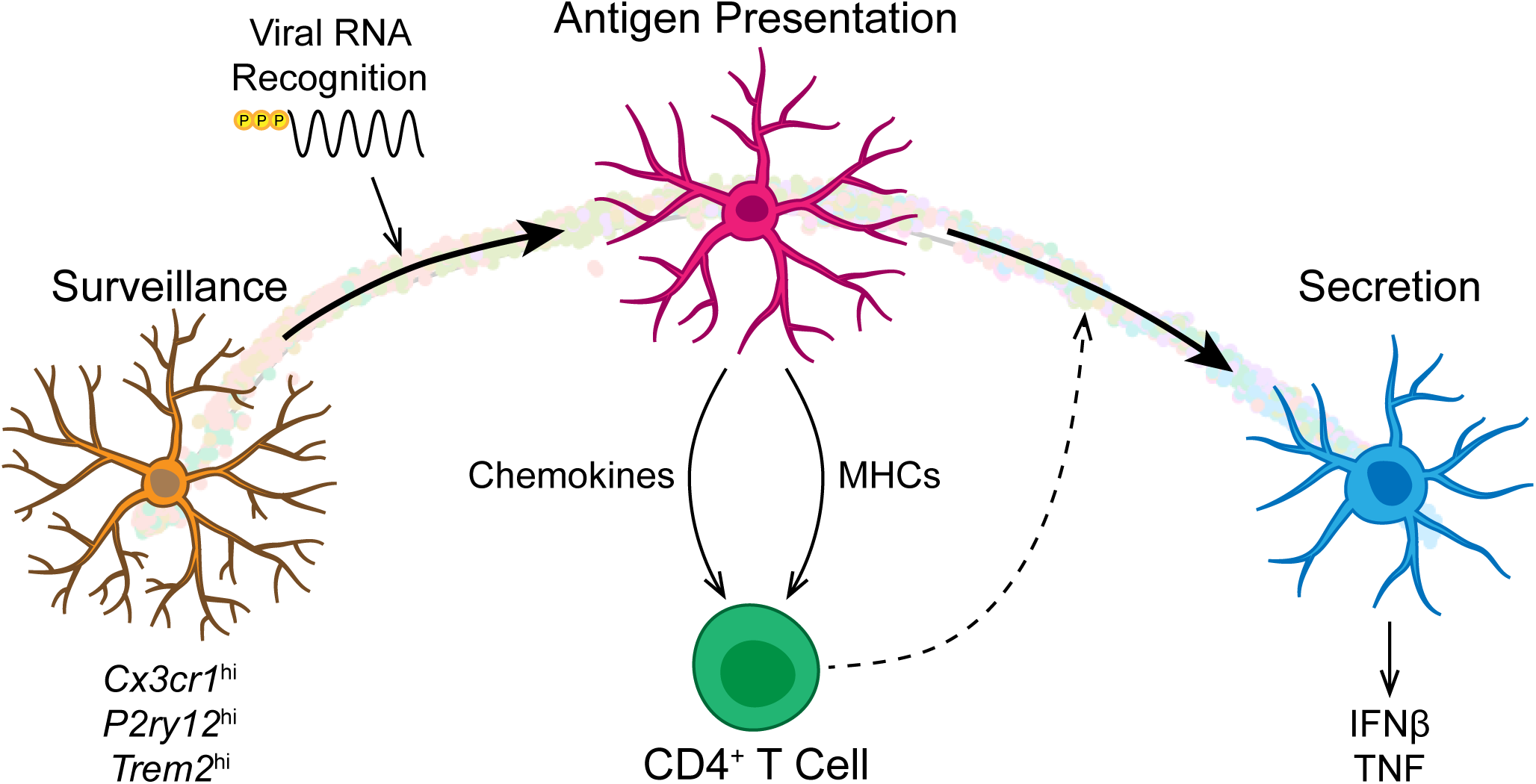
CD4^+^ T cells may facilitate microglial state transitions. Schematic of a model for microglial state transitions. Detection of viral RNAs induces the transition of microglia from a homeostatic surveillance state to an antigen presenting activated state, in which microglia up-regulate genes encoding chemokines and MHCs that subsequently activate infiltrating CD4^+^ T cells. Interactions between antigen presenting microglia and CD4^+^ T cells may facilitate the further progression of microglia along the activation trajectory with a transition into a secretory state with increased production of additional cytokines, including IFNβ and TNF.

An advantage of using scRNA-seq in this study has been the use of the transcriptome to assign and identify cells into distinct classes, types, and states. Despite the relatively low sequencing depth that we incur with the use of high-throughput droplet-based methods, our dataset captures variations across more dimensions than “conventional” techniques such as immunolabeling and *in situ* hybridization. The use of the transcriptome along with careful cross-referencing of gene expression signatures in cell clusters with well-validated studies also reduces biases in cell type identification or selection that may be introduced by the choice of markers. Consistent with other recent scRNA-seq studies, our results highlight potential biases introduced when using genes such as *Cx3cr1* for identifying microglia, since several of these genes are down-regulated in activated or disease-associated microglia^6–8, 43^.

In addition to resolving transcriptional differences between different immune cell types, our study also finds distinguishing gene expression features between distinct subclusters of microglia. While our results suggest that these subclusters represent discrete states along single activation trajectory, we do not rule out the possibility of distinct activation endpoints for microglia in a branching trajectory. It is possible that these differences may emerge when comparing across a variety of stimuli (e.g. “viral déjà vu”, LPS, AD transgenic models, experimental autoimmune encephalomyelitis) that may trigger different response pathways and a distinct set of activation trajectories from what we have observed in our study. Additionally, our study only examines transcriptional changes at a single time point after initial exposure to the virus, and may only capture a segment of microglial activation before the emergence of distinct endpoints. Future studies that systematically explore a full range of time points will help resolve the dynamics of these transcriptional responses. These studies will also clarify if the difference in leukocyte infiltration that we found to be correlated with the infection magnitude reflects a difference in the immune response, or a change in the time course of a shared response. Deeper sequencing and datasets containing a larger collection of microglia may also help resolve finer levels of heterogeneity among lower-expressing genes.

While our study describes transcriptional changes in CNS resident cells, it is also plausible that immune responses may differ across brain regions. These could be due to differences in the composition of resident cells, even among closely related types. For instance, resident MΦs in the DRN, a serotonergic nucleus, express the serotonin receptor *Htr1b*, which was not detected in resident MΦs in the cortex, striatum, or ventral midbrain^23, 44^. Interregional variation in immune signaling may also result from differences in the proximity of different locations to different neurovascular or ventricular features. The DRN is situated in close proximity to the cerebral aqueduct, as well as larger blood vessels in the ventrolateral periaqueductal gray that run along the anterior-posterior axis. Future studies comparing the responses in various regions (e.g. frontal cortex versus DRN) may reveal shared immune mechanisms across the CNS, and may also identify specific brain regions that are particularly susceptible to immunological insults that may subsequently trigger profound and long-lasting behavioral changes. Studies profiling these responses in different brain regions from the same animal may also distinguish systemic from local or region-specific effects. Although the injection sites in our study are over 1 mm away from the DRN, previous studies have shown increases in ISG expression in the cerebellum following infection of the olfactory bulb^45^. Since this study does not examine changes occurring at the injection sites or in regions devoid of RbV-infected neurons, we are unable to distinguish the effects of systemic signaling on the transcriptional responses that we describe here.

Many recent studies have used scRNA-seq to build detailed atlases of the diverse cell types that exist in the CNS. Relating the molecular profile of each cell type to its anatomical location and axonal projections remains as one of the main challenges in placing each of these cell types into neural circuits for functional studies, with the lower throughput of most anatomical tracing techniques being a major limiting factor. Methods for combining next-generation sequencing with connectivity mapping have therefore been of great interest, and several recently developed methods have used neurotropic viruses to either label cells for sorting and enrichment of transcripts from a projection-defined neuronal population^46, 47^, or the introduction of barcodes for reconstruction of neuronal connectivity from sequencing data^48, 49^. RbVs have been valuable tools in the study of neural connectivity since they can be used for cell type-specific transsynaptic retrograde tracing, which provides more specificity in the identity of the postsynaptic cell than conventional tracers^13^. While our study provides evidence further cautioning against the use of the SAD B19 strain for functional studies^50, 51^, we find that cells with high expression of RbV transcripts still retain sufficient transcriptional information for their classification into a specific cell type. Our results therefore support the feasibility of the use of RbVs for connectivity mapping between single cells using high-throughput sequencing methods. Although the detection of RbV-infected neurons was sparse in our dataset, limitations in the yield of RbV-labeled neurons for connectivity inference and network reconstruction can be overcome through the addition of methods for enriching for both RbV-labeled cells, such as FACS, or enrichment of RbV transcripts during library preparation. Future studies comparing the immune responses elicited by different viruses or virus strains used for neuroscience research will also provide crucial insights into the effects of these tools on the physiological properties of the cell types and neural circuits of interest.

## METHODS

### Mice

C57BL/6J (The Jackson Laboratory, Stock # 000664) were kept on a 12:12 regular light/dark cycle under standard housing conditions. All procedures were performed in accordance with protocols approved by the Harvard Standing Committee on Animal Care following guidelines described in the U.S. National Institutes of Health Guide for the Care and Use of Laboratory Animals (Harvard Medical School IACUC protocol #IS00000571).

### Rabies Viruses

Unpseudotyped rabies viruses (B19G-SADΔG-EGFP, B19G-SADΔG-tdTomato) were generated in-house using procedures based on published protocols^52, 53^. Virions were amplified from existing stocks in three rounds of low-MOI passaging through BHK-B19G cells by transfer of filtered supernatant, with 3 to 4 days between passages. Cells were grown at 35 °C and 5% CO_2_ in DMEM with GlutaMAX (Thermo Scientific, #10569010) supplemented with 5% heat-inactivated FBS (Thermo Scientific #10082147) and antibiotic-antimycotic (Thermo Scientific #15240-062). Virions were concentrated from media from dishes containing virion-generating cells by first collecting and incubating with benzonase nuclease (1:1000, Millipore #70664) at 37 °C for 30 min, followed by filtration through a 0.22 µm PES filter. The filtered supernatant was transferred to ultracentrifuge tubes (Beckman Coulter #344058) with 2 ml of a 20% sucrose in dPBS cushion and ultracentrifugated at 20,000 RPM (Beckman Coulter SW 32 Ti rotor) at 4 °C for 2 hours. The supernatant was discarded and the pellet was resuspended in dPBS for 6 hours on an orbital shaker at 4 °C before aliquots were prepared and frozen for long-term storage at -80 °C. Unpseudotyped rabies virus titers were estimated based on a serial dilution method counting infected HEK 293T cells, and quantified as infectious units per ml (IU/ml).

### Stereotaxic Surgeries

Mice were initially anesthetized with 5% isoflurane (80% oxygen) and maintained at 1-2.5% isoflurane after placement on the stereotaxic frame (David Kopf Instruments, Model 1900 Stereotaxic Alignment System). The scalp was cleaned and sterilized before an incision was made to expose the skull, and sterile ophthalmic ointment was applied to the eyes. For leveling the horizontal plane, a stereotaxic alignment tool (David Kopf Instruments, Model 1905) was used to zero the relative dorsoventral displacement of Bregma and Lambda, as defined in the Paxinos Brain Altas^54^, for adjusting tilt of the anterior-posterior axis, and of two points equidistant to the left and right of Bregma for adjusting the tilt of the medial-lateral axis. Craniotomies were prepared using a mounted drill (David Kopf Instruments, Model 1911) with careful removal of the bone flap and overlying dura using forceps and a fine needle tip, and were covered with sterile 0.9% saline before and during the injection to prevent desiccation. Viruses were front-filled into a pulled glass pipette (Drummond Scientific, #5-000-2005) filled with mineral oil (Millipore Sigma, M3516) and connected to a 5 µl Hamilton syringe (Hamilton #84850) via polyethylene tubing filled with mineral oil. Glass pipettes were pulled to obtain a tip size of approximately 40-60 µm on a pipette puller (Sutter Instrument Co., P-97). Viruses were infused into target regions at approximately 100 nl/min using a syringe pump (Harvard Apparatus, #883015), and pipettes were slowly withdrawn (< 10 µm/s) at least 10 min after the end of the infusion. Following wound closure, mice were placed in a cage with a heating pad until their activity was recovered before returning to their home cage. Mice were given pre- and post-operative oral carprofen (MediGel CPF, 5mg/kg/day) as an analgesic, and monitored daily for at least 4 days post-surgery.

### Stereotaxic Injection Coordinates and Volumes

All coordinates are relative to Bregma along the anterior-posterior axis and medial-lateral axis, and relative to the pial surface along the dorsoventral axis. “BL” denotes the distance between Bregma and Lambda. All injections used a straight vertical approach parallel to the DV (Z) axis. All injections were placed in the right hemisphere (positive ML values). Striatum (Str): AP = +0.40 mm, ML = ±2.45 mm, DV = -3.10 mm, 300 nl. Dorsal lateral geniculate nucleus (dLGN): AP = -(2.00 * BL / 4.20) mm, ML = +2.25 mm, DV = -3.00 mm, 150 nl. Substantia Nigra (SN): AP = -(3.00 * BL / 4.20) mm, ML = +1.32 mm, DV = -4.60 mm, 150 nl.

### Histology

Mice were deeply anesthetized with isoflurane and transcardially perfused with 5-10 ml chilled 0.1 M PBS, followed by 10-15 ml chilled 4% paraformaldehyde in 0.1 M PBS. Brains were dissected out and post-fixed overnight at 4 °C, followed by incubation in a storing/cryoprotectant solution of 30% sucrose and 0.05% sodium azide in 0.1 M PBS for at least 1-2 days to equilibrate. 50 µm coronal slices were prepared on a freezing microtome (Leica Biosystems, SM2010 R). 50 µm thick free-floating tissue sections were rinsed 3 x 5 min with 0.1 M PBS containing 0.5% Triton X-100 (PBST) before counterstaining with Neurotrace 435 (ThermoFisher Scientific N21479) at a concentration of 1:100 in 0.1 M PBS with 0.5% Triton X-100 for 1 hour at room temperature. Slices were rinsed 4 x 5 min with 0.1 M PBS before they were mounted on glass slides in VectaShield mounting media (Vector Labs, H-1000). Fluorescence images were taken on an Olympus VS120 slide scanning microscope with a 10X air objective.

### Single Cell Dissociation and RNA Sequencing

8- to 10-week old C57BL/6J mice were pair-housed in a regular 12:12 light/dark cycle room prior to tissue collection. Mice were transcardially perfused with an ice-cold choline cutting solution containing neuronal activity blockers (110 mM choline chloride, 25 mM sodium bicarbonate, 12 mM D-glucose, 11.6 mM sodium L-ascorbate, 10 mM HEPES, 7.5 mM magnesium chloride, 3.1 mM sodium pyruvate, 2.5 mM potassium chloride, 1.25 mM sodium phosphate monobasic, 10 µM (R)-CPP, 1 µM tetrodotoxin, saturated with bubbling 95% oxygen/5% carbon dioxide, pH adjusted to 7.4 using sodium hydroxide). Brains were rapidly dissected out and sliced into 250 µm thick coronal sections on a vibratome (Leica VT1000) with a chilled cutting chamber filled with choline cutting solution. Coronal slices containing the dorsal raphe were then transferred to a chilled dissection dish containing a choline-based cutting solution for microdissection. Dissected tissue chunks were transferred to cold HBSS-based dissociation media (Thermo Fisher Scientific Cat. # 14170112, supplemented to final content concentrations: 138 mM sodium chloride, 11 mM D-glucose, 10 mM HEPES, 5.33 mM potassium chloride, 4.17 mM sodium bicarbonate, 2.12 mM magnesium chloride, 0.9 mM kynurenic acid, 0.441 mM potassium phosphate monobasic, 0.338 mM sodium phosphate monobasic, 10 µM (R)-CPP, 1 µM tetrodotoxin, saturated with bubbling 95% oxygen/5% carbon dioxide, pH adjusted to 7.35 using sodium hydroxide) supplemented with an additional inhibitor cocktail (10 µM triptolide, 5 µg/ml actinomycin D, 30 µg/ml anisomycin) and kept on ice until dissections were completed. The remaining tissue was fixed in 4% paraformaldehyde in phosphate-buffered saline for histological verification. Dissected tissue chunks for each sample were pooled into a single tube for the subsequent dissociation steps. Tissue chunks were first mixed with a digestion cocktail (dissociation media, supplemented to working concentrations: 20 U/ml papain, 1 mg/ml pronase, 0.05 mg/mL DNAse I, 10 µM triptolide, 5 µg/ml actinomycin D, 30 µg/ml anisomycin) and incubated at 34 °C for 90 min with gentle rocking. The digestion was quenched by adding dissociation media supplemented with 0.2% BSA and 10 mg/ml ovomucoid inhibitor (Worthington Cat. # LK003128), and samples were kept chilled for the rest of the dissociation procedure. Digested tissue was collected by brief centrifugation (5 min, 300 *g*), re-suspended in dissociation media supplemented with 0.2% BSA, 1 mg/ml ovomucoid inhibitor, and 0.05 mg/mL DNAse I. Tissue chunks were then mechanically triturated using fine-tip plastic micropipette tips of progressively decreasing size. The triturated cell suspension was filtered in two stages using a 70 µm cell strainer (Miltenyi Biotec Cat # 130-098-462) and 40 µm pipette tip filter (Bel-Art Cat. # H136800040) and washed in two repeated centrifugation (5 min, 300 *g*) and re-suspension steps to remove debris before a final re-suspension in dissociation media containing 0.04% BSA and 15% OptiPrep (Sigma D1556). Cell density was calculated based on hemocytometer counts and adjusted to approximately 100,000 cells/ml. Single-cell encapsulation and RNA capture on the InDrop platform was performed at the Harvard Medical School ICCB Single Cell Core using v3 hydrogels based on previously described protocols ^20^. Suspensions were kept chilled and gently agitated until the cells were flowed into the microfluidic device. Libraries were prepared and indexed following the protocols referenced above, and sequencing-ready libraries were stored at -80 °C. Libraries were pooled and sequenced on an Illumina NextSeq 500 (High Output v2 kits).

### Sequencing Data Processing

NGS data was processed using previously a published pipeline in Python available at [https://github.com/indrops/indrops]19. Briefly, reads were filtered by expected structure and sorted by the corresponding library index. Valid reads were then demultiplexed and sorted by cell barcodes. Cell barcodes containing fewer than 250 total reads were discarded, and remaining reads were aligned to a reference mouse transcriptome (Ensembl GRCm38 release 87) using Bowtie 1.2.2 (m = 200, n = 1, l = 15, e = 100). For alignment, the mouse transcriptome was modified with the addition of genes from the SAD B19 rabies viruses and transgenes (*B19N, B19P, B19M, B19L, EGFP, tdTomato, AmCyan1*). Aligned reads were then quantified as UMI-filtered mapped read (UMIFM) counts. UMIFM counts and quantification metrics for each cell were combined into a single file sorted by library and exported as a gunzipped TSV file.

### Pre-Clustering Filtering and Normalization

Analysis of the processed NGS data was performed in R version 3.4.4 using the *Seurat* package version 2.3.1^22, 55^. A custom R script was used to combine the expression data and metadata from all libraries corresponding to a single batch, and cells with fewer than 500 UMIFM counts were removed. The expression data matrix (Genes x Cells) was filtered to retain genes with > 5 UMIFM counts, and then loaded into a *Seurat* object along with the library metadata for downstream processing. The percentage of mitochondrial transcripts for each cell (*percent.mito*) was calculated and added as metadata to the *Seurat* object. Cells were further filtered prior to dimensionality reduction (*Reads – min. 20,000, max. Inf; nUMI – min. 500, max. 18,000; nGene – min. 200, max. 6,000; percent.mito – min. -Inf, max. 0.1*). Low quality libraries identified as outliers on scatter plots of quality control metrics (e.g. unusually low gradient on the nGene vs. nUMI) were also removed from the dataset. Expression values were then scaled to 10,000 transcripts per cell and log-transformed. Effects of latent variables (*nUMI, percent.mito, Sex*) were estimated and regressed out using a GLM (*ScaleData* function, *model.use* = *“linear”*), and the scaled and centered residuals were used for dimensionality reduction and clustering.

### Dimensionality Reduction and Batch Effect Correction

Canonical correlation analysis (CCA) was used for dimensionality reduction and mitigation of batch effects. We used 2,412 genes that were highly variable in at least 2 datasets to calculate canonical variates (CVs) using the *RunMultiCCA* function in *Seurat*. After inspection of the CVs, the first 21 CVs were used for subspace alignment using the *AlignSubspace* function to merge datasets into a single object.

### Cell Clustering and Cluster Identification

Initial clustering was performed on the merged and CCA-aligned dataset using the first 21 aligned CVs. UMAP was used only for data visualization. Clustering was run using the *FindClusters* function using the SLM algorithm and 10 iterations. Clustering was performed at varying resolution values, and we chose a value of 2 for the resolution parameter for the initial stage of clustering. Clusters were assigned preliminary identities based on expression of combinations of known marker genes for major cell classes and types. Low quality cells were identified based on a combination of low gene counts, low UMIFM counts, high fraction transcripts from mitochondrial genes, and a high fraction of nuclear transcripts (e.g. *Malat1*, *Meg3*, *Kcnq1ot1*). These cells typically clustered together and were removed manually. Following assignment of preliminary identities, cells were divided into data subsets as separate *Seurat* objects (neurons; astrocytes; ependymal cells; endothelial cells, pericytes, fibroblasts, and myocytes; immune cells; oligodendrocytes and polydendrocytes) for further subclustering.

### Subclustering

Subclustering was performed iteratively on each data subset to resolve additional cell types and subtypes. For immune cell types with proliferating cell populations (microglia, lymphocytes), cell cycle scores were calculated and regressed out using the *ScaleData* function in Seurat. Briefly, clustering was run at high resolution, and the resulting clusters were ordered in a cluster dendrogram built using the *Ward2* method in *hclust* using cluster-averaged gene expression for calculating the Euclidean distance matrix. Putative doublets/multiplets were identified based on expression of known marker genes for different cell types not in the cell subset (e.g. neuronal and glial markers). Putative doublets tended to separate from other cells and cluster together, and these clusters were removed from the dataset. Cluster separation was evaluated using the *AssessNodes* function and inspection of differentially expressed genes at each node. Clusters with poor separation, based on high OOBE scores and differential expression of mostly housekeeping genes, were merged to avoid over-separation of the data. The dendrogram was reconstructed after merging or removal of clusters, and the process of inspecting and merging or removing clusters was repeated until all resulting clusters could be distinguished based on a set of differentially expressed genes that we could validate separately.

### Differential Expression Tests and Gene Set Enrichment Analysis (GSEA)

Tests for differential expression (DE) were performed using *MAST* version 1.4.1^56^. P values were corrected using the Benjamini-Hochberg method and filtered a 5% false discovery rate (Q < 0.05; *n* = 18,737 genes). GSEA was performed using the *fgsea* package version 1.4.1 in R^57^. Genes were ordered by Z scores from *MAST* DE tests on either the *MSigDB* mouse Hallmark gene sets or Reactome pathways separately. Combined Z scores were used for most genes, and discrete component Z scores were used for genes in which the continuous component was returned as *NA* values (e.g. gene was not expressed in one of the two comparison groups). Enrichment scores were calculated using *fgsea* (*nperm = 100,000*, *maxSize* = *Inf*). P values were corrected in *fgsea* using the Benjamini-Hochberg method. Gene sets and pathways were obtained using the *misgdbr* package version 6.2.1.

### Trajectory Inference

Trajectory inference was performed using *monocle* version 2.6.4^39, 40^. Raw count data from the *Seurat* microglia object was converted to a *CellDataSet* object using the *importCDS* function in *monocle*. Genes that were differentially expressed between microglial subclusters (Q value < 0.01; |average log_2_ fold change| ≥ 1) were set as the ordering genes. The minimum spanning tree was constructed using the *reduceDimensions* function (*reduction_method = “DDRTree”*; *num_dim* = 10; *norm_method = “log”*; *residualModelFormula = “∼BatchID + nUMI + percent.mito”*; *relative_expr = TRUE; scaling = TRUE*).

### Inference of Intercellular Interactions

Intercellular interactions were inferred using the *CellPhoneDB* version 2.0 package in Python^41, 42^. A custom R script was used to export the single cell gene expression data from the curated *Seurat* object into a counts text file and metadata text file as recommended by the developers. Only genes with human orthologs were used, and mouse gene symbols were converted to the human ortholog gene symbols before data export using data from the *e!Ensembl* web portal. Data was processed in *CellPhoneDB* with statistical analysis (default parameters; iterations = 1,000; no sub-sampling). Data visualizations were made using *pheatmap* and *ggplot2* in R based on the plotting functions provided in the *CellPhoneDB* package.

### Statistics

Statistical tests and corrections for multiple testing were performed through the respective analysis packages described in each Methods section.

## Materials and Correspondence

Further information and requests for reagents may be directed to, and will be fulfilled by, the corresponding author Bernardo L. Sabatini.

## Data Availability

Sequencing data from rabies-injected animals generated in this study is available at NCBI GEO (accession number: GSE136455). The control dataset from uninjected animals was described in an earlier publication^23^, and is available at NCBI GEO (accession number: GSE134163).

## Code Availability

No custom packages were developed for the analysis in this study. Code documenting the data analysis in the form of R markdown notebooks are available upon request.

## ACKNOWLEDGEMENTS

We thank M. Hyun for assistance with tissue collection and scRNA-seq library preparation, N.E. Ochandarena for assistance with stereotaxic surgeries, and A.C. Philson for assistance with rabies virus production. We also thank the HMS ICCB Single Cell Core for assistance with scRNA-seq experiments on the InDrop platform; S. Hrvatin and A. Nagy (Greenberg Lab, HMS) for advice and help with scRNA-seq protocols and analysis pipelines; B.K. Lim (UCSD) and I.R. Wickersham (MIT) for advice and reagents for rabies virus production; the Bauer Core Facility at Harvard University for sequencing support; the HMS Neurobiology Imaging Facility for confocal microscopy support (P30 NS072030); J. Levasseur for assistance with animal husbandry; and L. Worth for administrative assistance; A. Saunders (HMS) for feedback on our manuscript; and members of the Sabatini Lab for helpful discussions. This work was supported by funding from the Howard Hughes Medical Institute (B.L.S.), National Institutes of Health (R01 MH100568 and R01 NS103226 to B.L.S.), a Harvard Brain Initiative Bipolar Disorder Seed Grant (B.L.S.), the HMS Department of Neurobiology Graduate Fellowship (K.W.H.), and the HMS Stuart H.Q. & Victoria Quan Fellowship in Neurobiology (K.W.H.).

## AUTHOR CONTRIBUTIONS

K.W.H. performed the experiments and data analysis. K.W.H and B.L.S. designed the experiments and wrote the manuscript.

## SUPPLEMENTARY INFORMATION

**Supplementary Fig. 1:**
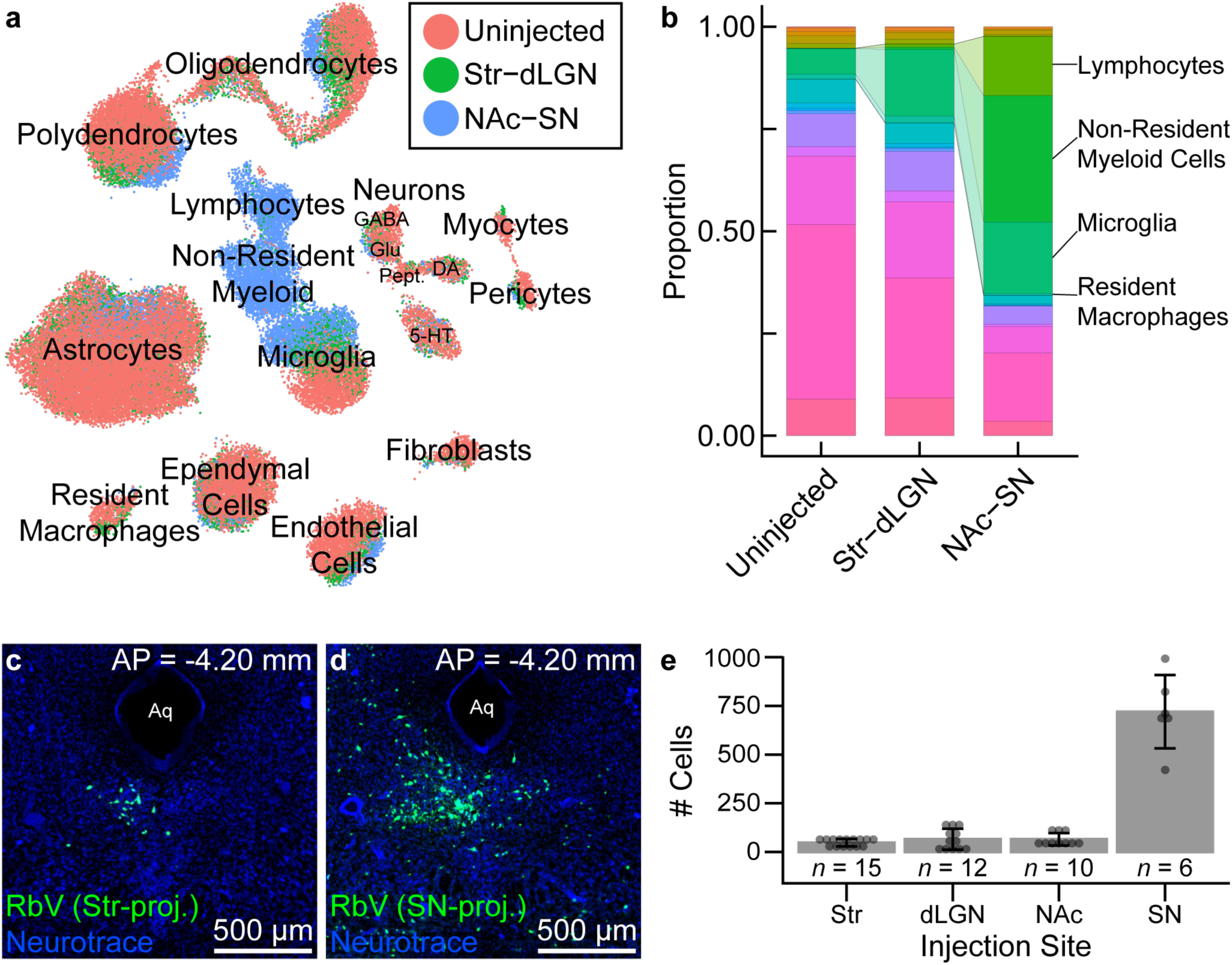
Leukocyte recruitment varies with infection magnitude. **(a)** UMAP plot of merged dataset with each cell color-coded by the injection site grouping. Specific neuron types are indicated and labeled by their primary neurotransmitter. **(b)** Stacked bar plot showing the relative proportion of each cell class/type in each of the injection site groups. **(c–d)** Fluorescence image of a representative coronal section showing neurons in the DRN and surrounding ventrolateral periaqueductal gray infected by RbV injected into the ipsilateral Str (**c**) or SN (**d**). All injections sites were targeted in the right hemisphere, and AP distances are relative to Bregma with negative values indicating positions posterior to Bregma. Injection volumes and virus titers were matched across injection sites. Abbreviations: Aq – cerebral aqueduct. dLGN – dorsal lateral geniculate nucleus. NAc – nucleus accumbens. SN – substantia nigra. Str – striatum. Scale bars: 500 µm. All coronal sections were counterstained with Neurotrace 435. **(e)** Bar plots showing the mean number RbV labeled cells for different injection sites. Counts from individual mice are represented as points, and the height of each bar indicates the mean for each group. Error bars are S.E.M. and *n* indicates the number of mice per group.

**Supplementary Fig. 2:**
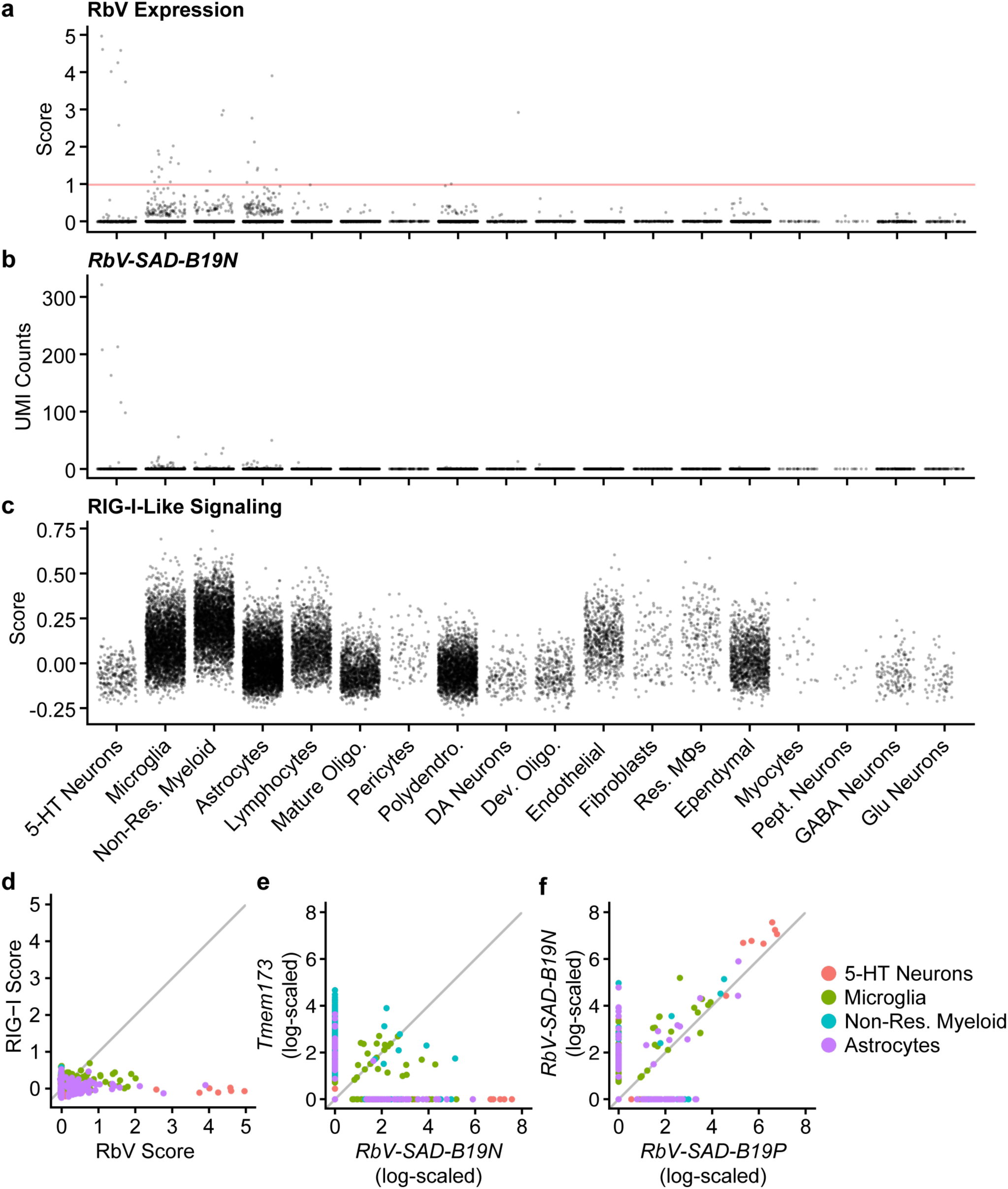
RbV transcripts are detected in neurons and phagocytic cells. **(a–c)** Dot plots showing the RbV gene set expression scores (**a**), *RbV-SAD-B19N* transcript counts (**b**), and RIG-I-Like signaling gene set expression scores (**c**) for individual cells grouped by cell class/type. Only cells from the RbV group are displayed. Cell classes/types (columns) are ordered horizontally by the average RbV expression score. 5-HT neurons have the highest RbV expression scores, followed by microglia, non-resident myeloid cells, and astrocytes. Score threshold is denoted by the horizontal red line. **(d–f)** Scatter plots for gene set expression scores (**d**) and log-scaled gene expression (**e**, **f**) in cells from the four cell types with the highest average RbV gene set expression scores. Points are single cells color coded by cell. No correlation was observed between the RIG-I-like signaling gene set expression score and RbV gene set expression score (**d**). Expression of *Tmem173* (STING) was highest in non-resident myeloid and microglia, but was low in 5-HT neurons even in cells with high expression of the *RbV-SAD-B19N* (**e**). Expression of genes within the RbV gene set, such as *RbV-SAD-B19N* and *RbV-SAD-B19P* were correlated (**f**). *RbV-SAD-B19N* expression was also higher compared to all other RbV genes, as previously described. Gray lines in D-F indicate the line of unity (*x* = *y*).

**Supplementary Fig. 3:**
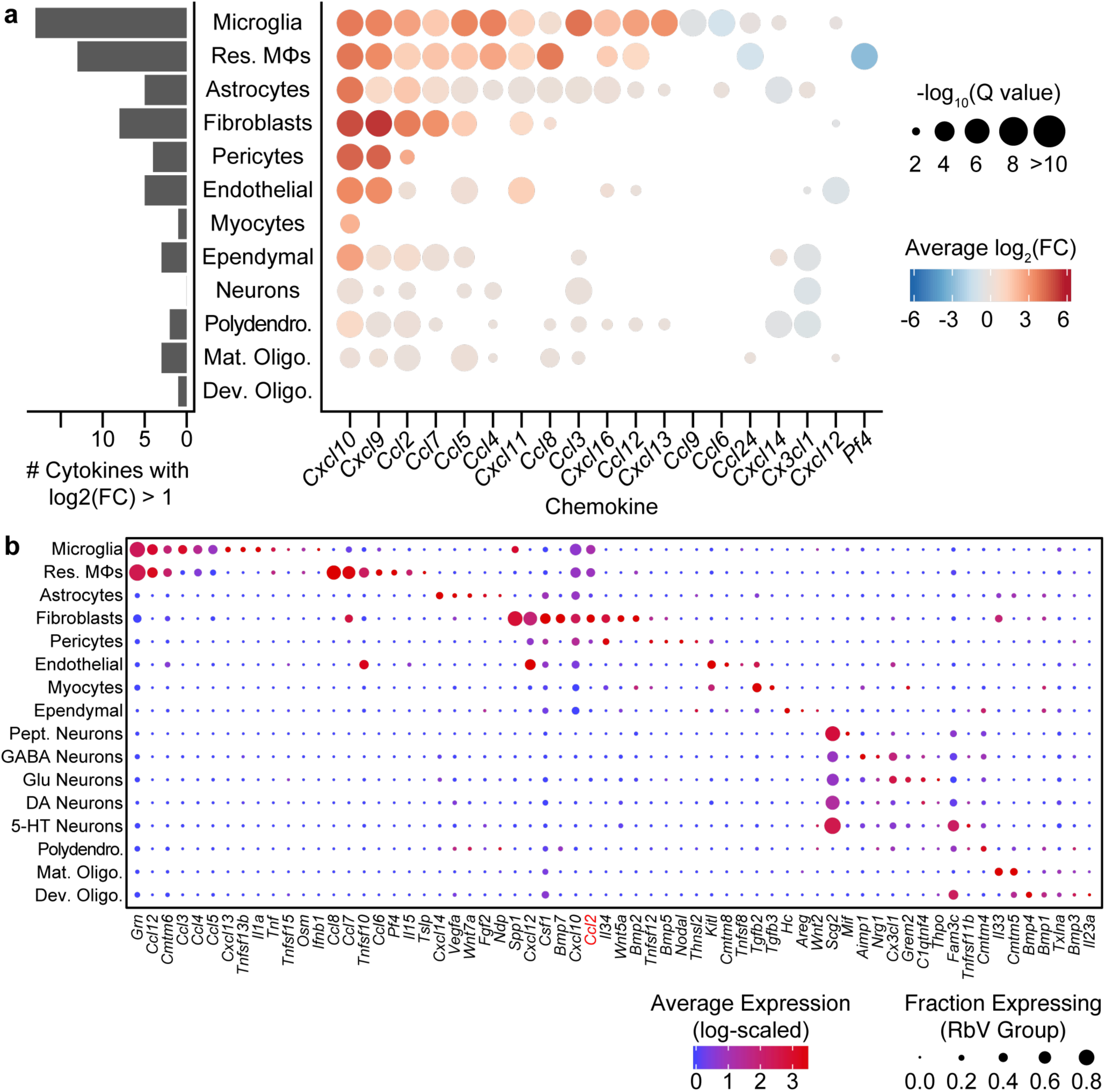
Cell type-specific expression of cytokines. **(a)** *Left*, Bar plot showing the number of significantly up-regulated cytokines (average log_2_ fold-change > 1) in each resident cell type. *Right,* Dot plot showing the cell type-specific changes in differentially expressed chemokines. The color of each dot represents the average log_2_ fold-change for each cell type cluster, and the size of the dot is scaled by the negative log_10_ of the adjusted P values (Q values, Benjamini-Hochberg correction). Points in which a gene was not considered differentially expressed for a given cell type (Q value ≥ 0.01) are not displayed. **(b)** Dot plot showing expression of selected cytokine genes (columns) by cell type (rows). The color of each dot represents the average log-scaled expression of each gene in a given cell type, and the size of each dot represents the fraction of cells of that type in which transcripts for that gene were detected.

**Supplementary Fig. 4:**
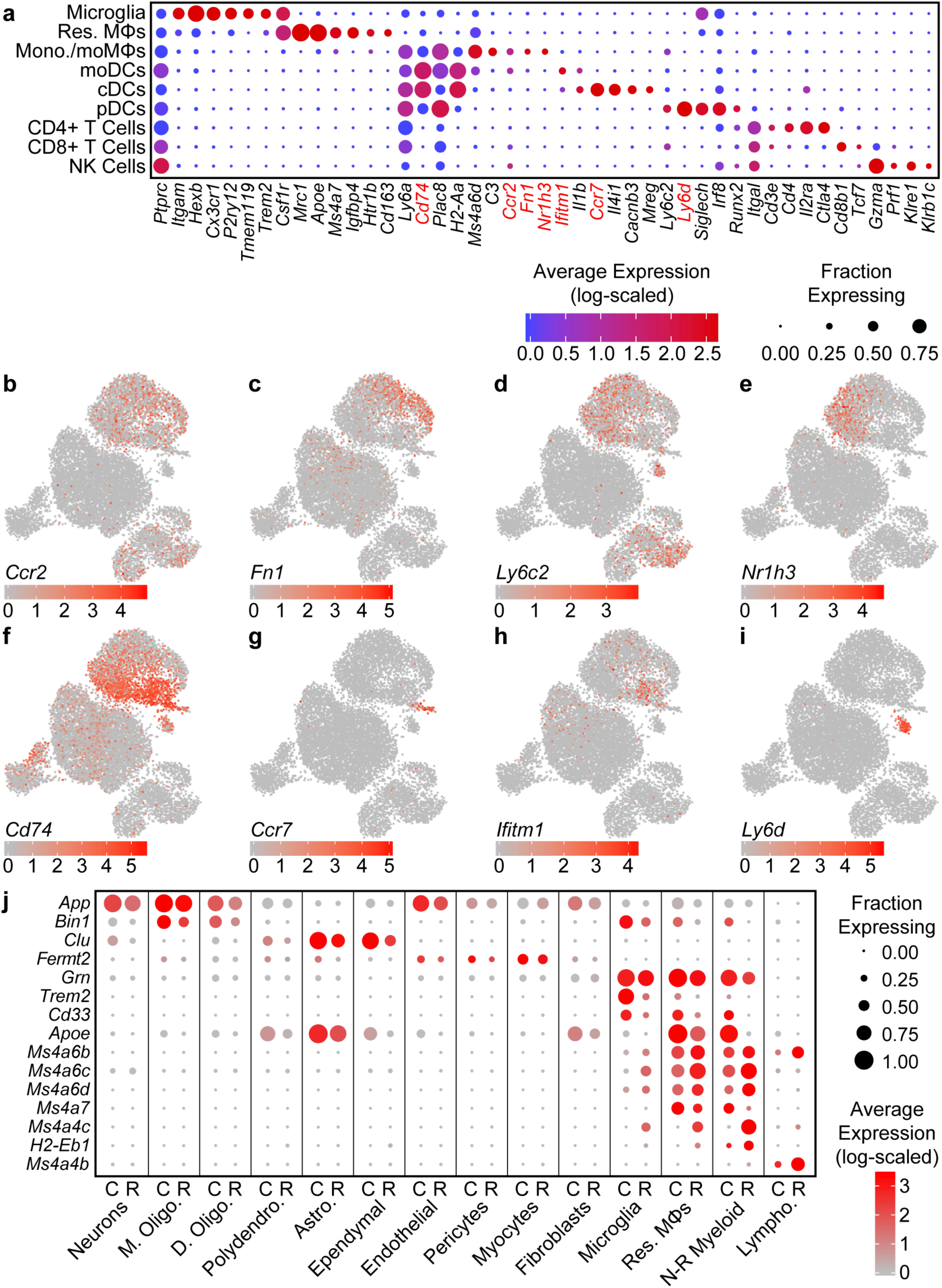
Identification of immune cell clusters. **(a)** Dot plot showing expression of several example genes (rows) used for identifying immune cell clusters (columns). **(b–i)** UMAP feature plots showing the expression of genes that are differentially expressed between non-resident myeloid cell types. Each cell is color-coded by its log-scaled expression for the indicated gene. **(j)** Paired dot plot showing expression of several genes associated with Alzheimer’s Disease (rows), separating by cell type (columns) and condition (sub-columns: C – Control; R – RbV).

**Supplementary Fig. 5:**
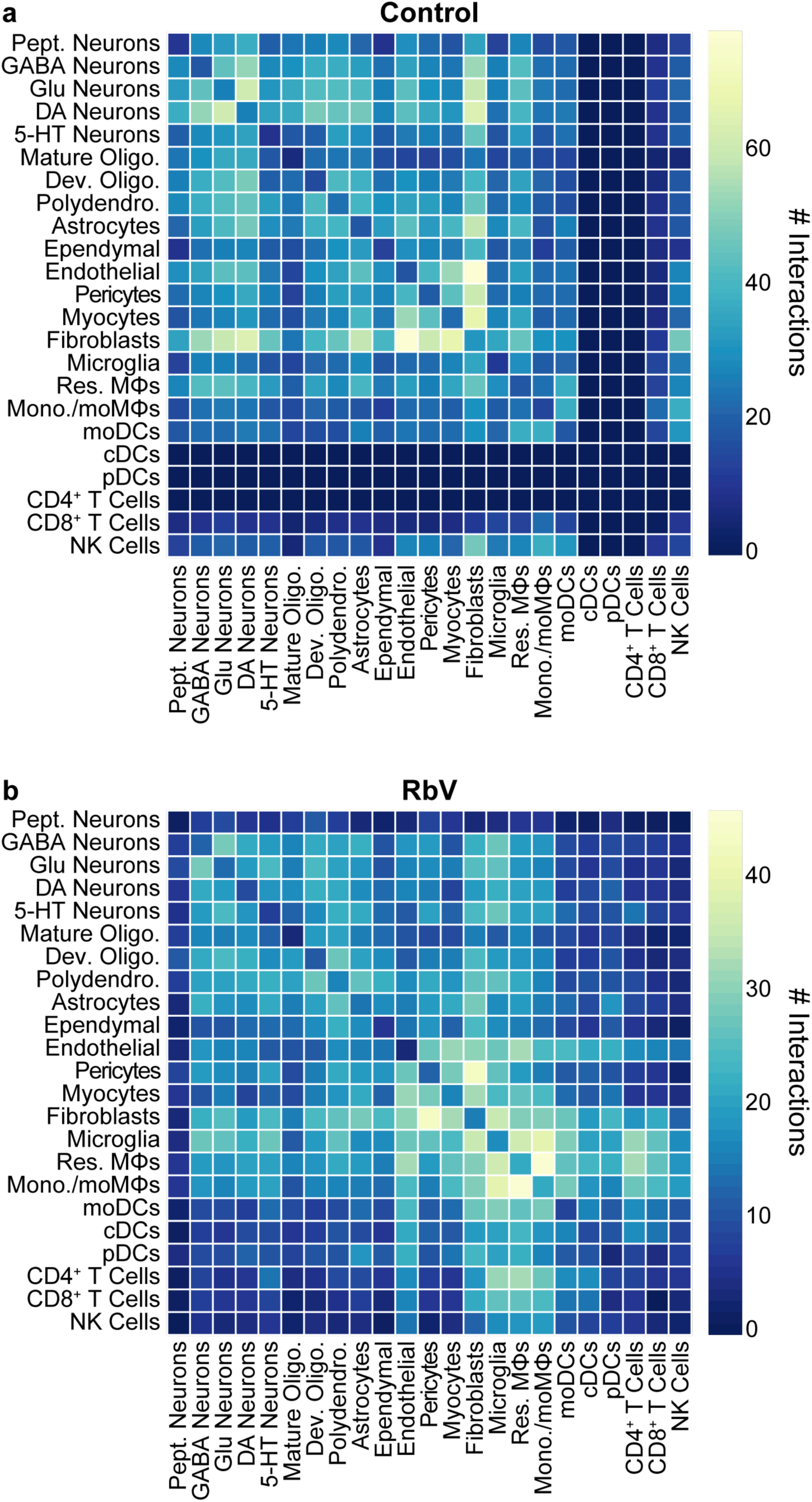
Number of inferred interactions between cell types by condition. **(a)** Heatmap showing the number of inferred interactions between cell types in the control group. The color range is scaled to the maximum number of interactions between a pair of cell types in the Control group. Interaction counts are non-directional, and not weighted by cell type abundance. Interactions for cell types not present in the Control group (e.g. dendritic cells) are at 0. **(b)** Heatmap showing the number of inferred interactions between cell types in the RbV group. The color range is scaled to the maximum number of interactions between a pair of cell types in the RbV group.

